# A biochemical probe for microtubule lattice integrity uncovers motor-caused lattice damage

**DOI:** 10.64898/2026.03.16.711994

**Authors:** Cornelia Egoldt, Luc Reymond, Joshua Tran, Aleksandar Salim, Marie-Claire Velluz, Sascha Hoogendoorn, Charlotte Aumeier

**Affiliations:** Department of Biochemistry, University of Geneva, 1211 Geneva, Switzerland; Biomolecular Screening Facility, EPFL, 1015 Lausanne, Switzerland; Department of Organic Chemistry, University of Geneva, 1211 Geneva, Switzerland

**Keywords:** Microtubule damage, in vitro damage probe

## Abstract

Microtubules experience mechanical and enzymatic stresses that can compromise lattice integrity, yet where and how lattice damage forms remains poorly understood due to the lack of tools that directly visualize damage as it occurs. Existing approaches infer damage indirectly through repair events or rely on static ultrastructural snapshots, precluding dynamic analysis. Here, we introduce MT-DS (Microtubule Damage Sensor), a damage-selective fluorescent probe that directly labels microtubule lattice openings. MT-DS combines taxane-based microtubule binding with a multivalent protein scaffold to restrict intraluminal diffusion and selectively retain the probe at sites of lattice openings. Using MT-DS, we visualize intrinsic lattice defects in stabilized microtubules, uncover a strong enrichment of damage at annealing sites, and demonstrate that kinesin-1Δ6 actively generates de novo lattice damage during motility. By enabling direct, time-resolved detection of microtubule damage, MT-DS establishes lattice integrity as an experimentally accessible parameter and provides a chemical tool to investigate how mechanical stress reshapes the microtubule cytoskeleton.

## INTRODUCTION

Microtubules are dynamic cytoskeletal polymers that organize intracellular transport, cell division, and cell polarity^1,2^. While their dynamic instability has been studied in detail, the structural integrity of the microtubule lattice itself has only recently emerged as a regulatory layer of microtubule function^3^. During their lifetime, microtubules are exposed to mechanical strain, enzymatic severing, and forces generated by molecular motors that use the lattice as a track^4–6^. These stresses can locally destabilize the lattice, increasing the probability of tubulin dimer dissociation from the microtubule shaft^7^.

Growing evidence indicates that lattice damage sites are not passive structural defects but functionally relevant elements of the microtubule cytoskeleton. In vitro and in cells, repaired damage sites act as rescue sites where depolymerizing microtubules halt shrinkage and resume growth, thereby increasing microtubule lifetime and length^3,8,9^. Through this mechanism, lattice damage and repair influence microtubule network organization and cell polarity^10^. In addition, damage sites selectively recruit proteins that facilitate repair, and repaired lattice regions display altered post-translational modification patterns, indicating that damage locally rewires the biochemical identity of the microtubule^5,9,11–14^. Together, these findings establish lattice damage as a dynamic and biologically meaningful feature of microtubule regulation. However, where damage sites form, how they arise, and how they evolve over time remain largely unknown.

This knowledge gap is primarily methodological. Microtubule lattice defects can arise during polymerization through protofilament number mismatches or geometric discontinuities, and in assembled lattices through spontaneous tubulin loss^15–17^. Defects can be repaired by incorporation of free GTP-tubulin, a process visualized in vitro using differentially labeled tubulin^3,9,18^. While these approaches demonstrated that self-repair occurs, they report only on repair events, provide limited temporal resolution, and do not allow direct detection of unrepaired or transient damage sites. Direct visualization of lattice damage has so far been restricted to electron microscopy of fixed samples, which provides structural detail but is low-throughput and incompatible with dynamic measurements^5,19,20^.

A particularly prominent and unresolved source of lattice stress is molecular motor activity. Kinesin-1 has been shown to destabilize the microtubule lattice, and the kinesin-1Δ6 mutant induces pronounced shaft damage during motility^7,19,21^. However, it remains debated whether motor proteins generate lattice damage de novo or primarily amplify pre-existing defects arising from imperfect polymerization. Resolving this question requires a tool that can directly and dynamically detect microtubule damage as it forms, a capability that has been missing from the cytoskeletal toolbox.

Here, we introduce MT-DS (Microtubule Damage Sensor), a damage-selective fluorescent probe that directly labels microtubule lattice openings. MT-DS combines taxane-based microtubule binding with a multivalent protein scaffold to reduce intraluminal diffusion and selectively retain the probe at lattice discontinuities. Unlike existing microtubule probes^22,23^, MT-DS does not label the intact lattice uniformly but instead reports sites of lattice openings with high spatial and temporal resolution.

Using MT-DS, we visualize intrinsic lattice defects in stabilized microtubules, uncover a strong enrichment of damage at annealing sites, and demonstrate that kinesin-1Δ6 actively generates de novo lattice damage while moving along microtubules. By enabling direct, real-time detection of microtubule damage, MT-DS establishes lattice integrity as an experimentally accessible parameter and provides a chemical tool to interrogate how mechanical stress reshapes the microtubule cytoskeleton.

## RESULTS

### Current approaches provide only indirect or static views of microtubule lattice damage

Microtubule lattice damage can be inferred in vitro using slow lattice disassembly assays of GMPCPP-stabilized microtubules, in absence of free tubulin, imaged by total internal reflection fluorescence (TIRF) microscopy. Under these conditions, lattice destabilization manifests in two characteristic ways: the formation of pronounced kinks that often precede microtubule breakage, and progressive lattice weakening, visible as longitudinal decreases in fluorescence intensity along the microtubule shaft. These weakened regions are mechanically unstable and frequently fluctuate in and out of the focal plane (Fig. 1a).

**Fig. 1.**
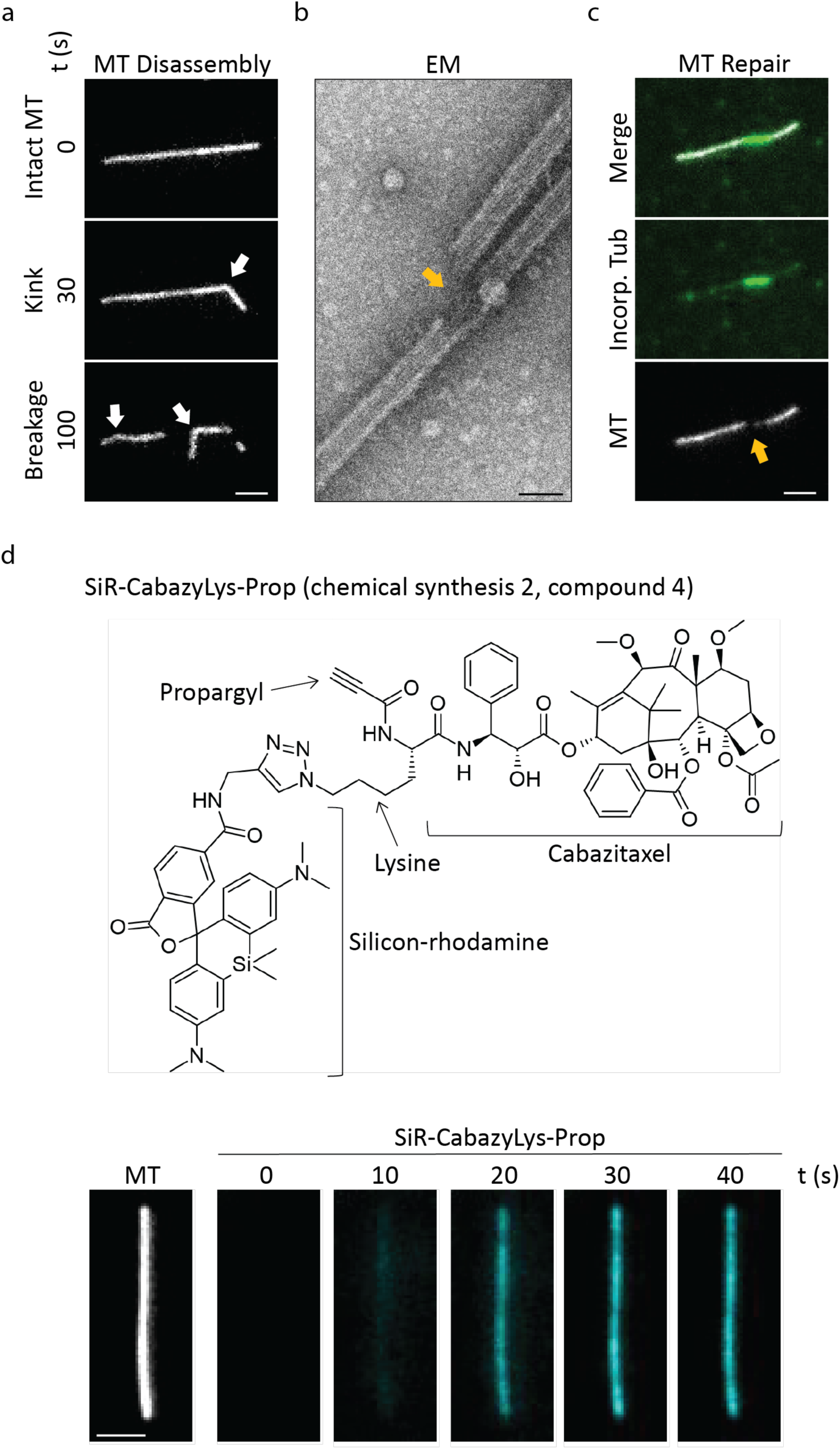
Limitations of microtubule damage detection in vitro. (**a**) Representative fluorescence images of a GMPCPP-stabilized microtubule (MT, white) undergoing kink formation after 30 s, followed by lattice softening leading to breakage after 100 s, in absence of free tubulin and anti-bleaching buffer, indicated by white arrows. For characterization and quantification of this effect see^6,30^. Scale bars: 2 μm. (**b**) Electron microscopy (EM) image of damaged microtubule. Damage - yellow arrow. Scale bar: 50nm. (**c**) Representative fluorescence images showing repair of lattice damage through incorporation of free tubulin (Incorp. Tub, green) into a GMPCPP-microtubule, gray). Scale bars: 2 μm. (**d**) Top, Chemical structure of the SiR-tubulin derivative SiR-CabazyLys-Prop, compound4. Bottom: time series showing microfluidic-based labeling of a GMPCPP-stabilized microtubule (white) upon addition of the SiR-CabazyLys-Prop (10nM, cyan). Scale bar: 2 μm.

While such assays report macroscopic lattice failure, they do not resolve the underlying structural defects. Direct visualization of lattice damage at molecular resolution has so far relied on electron microscopy (EM), which reveals local losses of tubulin density at damage sites (Fig. 1b). However, EM requires fixation and therefore captures only static snapshots, precluding analysis of damage formation, progression, and repair dynamics.

An alternative fluorescence-based strategy relies on the incorporation of free, fluorescently labeled tubulin into lattice defects (Fig. 1c). This repair assay has been instrumental in demonstrating microtubule self-repair but reports exclusively on repair events rather than damage itself. Moreover, high background from nonspecific tubulin adhesion, limited temporal resolution, and the need for long incubation times restrict its ability to capture dynamic or transient damage sites. Together, these limitations underscore the absence of a tool capable of directly and dynamically detecting microtubule lattice damage.

### Design and assembly of the microtubule damage sensor MT-DS

To overcome these limitations, we sought to develop a probe that selectively labels lattice openings while remaining excluded from intact microtubules. We chose a taxane-based microtubule binder as it specifically binds to the microtubule lumen, exemplified by SiR-tubulin which consists of a silicon-rhodamine (SiR)-conjugated derivative of docetaxel^22^. We synthesized SiR-CabazyLys-Prop (Compound 4), a cabazitaxel-derived taxane functionalized with a Lys-N_3_ linker to enable conjugation of alkyne-functionalized SiR via click chemistry (Fig. S1a and Extended Methods). However, like SiR-tubulin and taxol^24^, this small-molecule probe showed rapid intraluminal diffusion, preventing selective labeling of defined microtubule entry sites such as lattice defects or microtubule ends (Fig. 1d). These observations motivated the design of a probe that retains taxane-based microtubule binding while markedly reducing diffusion within the lumen.

As an initial strategy to increase probe size, we conjugated SiR-CabazyLys-Prop to dextran scaffolds of two different sizes (40 or 70 kDa). The propargyl group of SiR-CabazyLys-Prop enabled multivalent coupling by reacting with PEG₃–N₃–functionalized amino-dextran using click chemistry, generating two size-distinct dextran-based probes 8a and 8b, carrying approximately seven taxane–fluorophore units (Fig. S1b and Extended Methods). While this approach successfully increased probe size, the resulting probe strongly self-aggregated and showed low diffusive labeling along the microtubule instead of localized damage detection (Fig. S1c).

We therefore pursued an alternative design based on a defined, self-assembling protein scaffold. MT-DS consists of a multivalent protein scaffold decorated with multiple taxane-based microtubule binders and fluorophores. As a scaffold, we used a previously engineered 24-subunit self-assembling protein nanostructure^25^, onto which HaloTag7 domains were genetically fused (Fig. 2). This design enables controlled, 24-valent functionalization while increasing the effective size of the probe.

**Fig. 2.**
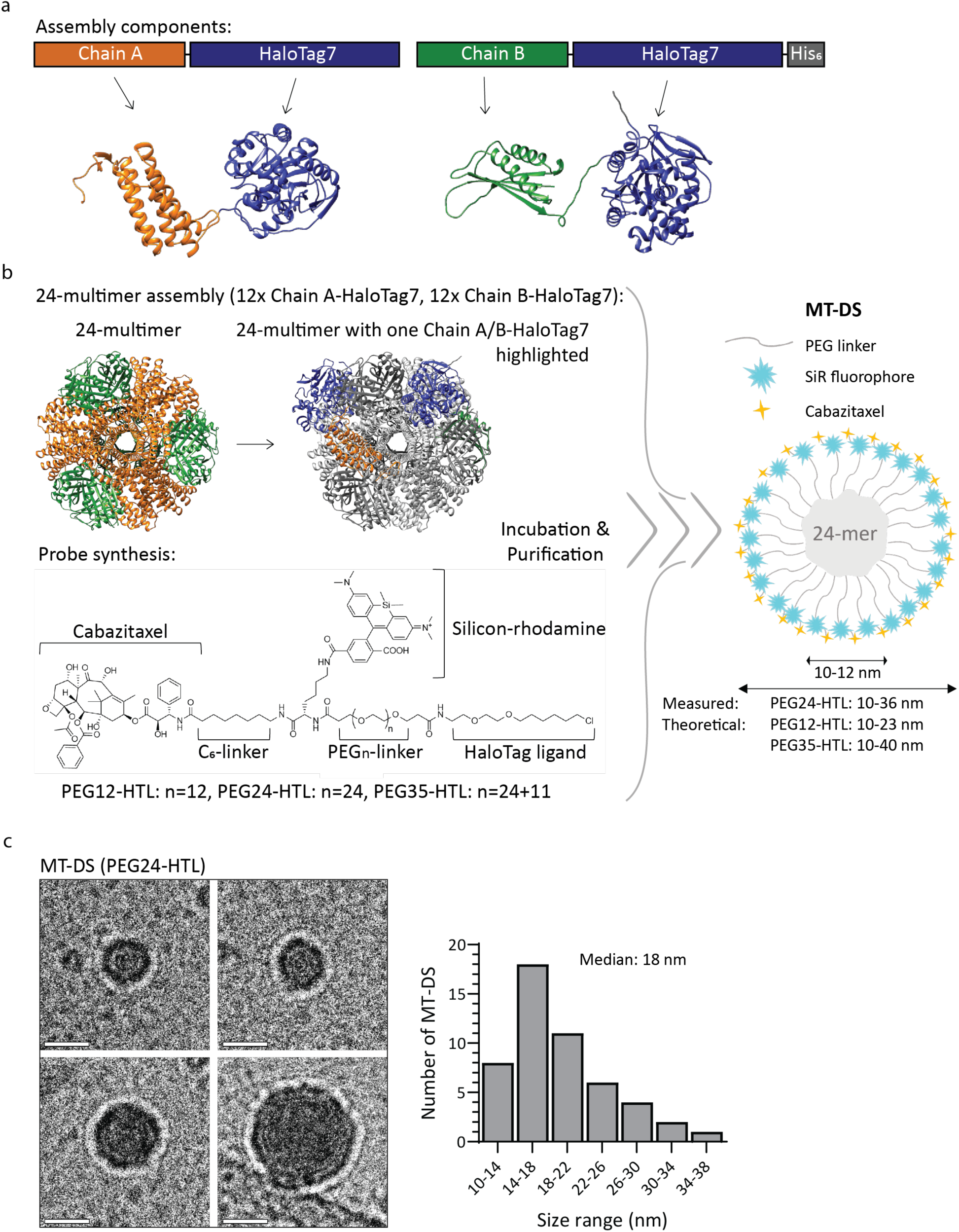
Generation and characterization of the microtubule damage sensor MT-DS. (**a**) Scheme displaying assembly components of the 24-mer Chain A-HaloTag7 and Chain B-HaloTag7-H (His_6_-tag). Predicted structures of each subunit generated with AlphaFold3 are shown below^39^. (**b**) Assembly principle of MT-DS. MT-DS consists of two independent components - the protein multimer scaffold and the chemical probe. The protein scaffold consists of two distinct protein components, Chain A (orange) and Chain B (green), that self-assemble into a 24-subunit cage-like complex (24-mer) corresponding to the architecture described in PDB: 4NWP, according to King et al.^25^. Each component forms a trimer, and four copies of Chain A trimer and Chain B trimer each assemble via designed interfaces to form the nanocage. To illustrate the impact of the additional HaloTag7 (blue), one Chain A-HaloTag7 and one Chain B-HaloTag7 protein is projected on the 24-mer structure. The chemical probe design is derived from SiR-tubulin^21^ and modified with a PEG_n_-linker (PEG12-HTL, PEG24-HTL, PEG35-HTL) and chloroalkane HaloTag-ligand. After purification of the 24-mer scaffold and synthesis of the probe, both components are combined to generate MT-DS variants with different linker lengths (PEG12-HTL, PEG24-HTL, and PEG35-HTL; see Methods and Fig. S1). Theoretical particle diameters range from 10 to 40 nm. (**c**) Cryo-electron microscopy images of PEG24-HTL particles. Observed particle diameters range from 10 to 38 nm. Scale bar, 15 nm. Right, size distribution of MT-DS particles (n = 50).

To generate MT-DS, we synthesized HaloTag ligands containing a SiR fluorophore and a taxane derivative connected via polyethylene glycol (PEG) linkers of defined length. By varying PEG length and ligand composition, we produced a series of MT-DS variants with tunable dimensions and microtubule-binding valency, yielding progressively larger probes with increasing linker length (PEG12-HTL 11, PEG24-HTL 10, PEG35-HTL 12; Fig. 2b and Fig. S1a and Extended Methods). The resulting architecture comprises a 24-subunit protein nanosphere decorated with flexible PEG arms bearing the fluorophore and microtubule-binding moiety ranging in size from 10 nm to 38 nm for PEG24-HTL (Fig. 2c). This design enables multivalent interaction with lattice openings while restricting intraluminal diffusion. To further modulate affinity without altering probe size, we partially substituted PEG24-HTL with a non-microtubule-binding HaloTag fluorophore (SPY555-CA), generating a MT-DS variant with approximately 50% reduced binding valency (Fig. S2b).

Following assembly and purification by size-exclusion chromatography (SEC), all MT-DS variants formed monodisperse complexes with complete HaloTag labeling (Fig. 2). After SEC purification, this design yields a pure and stable assembled probe that can be used directly or stored at –80°C upon usage (Fig. 2c, Methods).

### MT-DS selectively labels microtubule ends and distinct lattice sites

We next evaluated MT-DS performance on GMPCPP-stabilized microtubules. Because microtubule ends expose luminal binding sites, they provide a straightforward initial test of probe functionality. All MT-DS variants rapidly labeled microtubule ends. In addition, discrete puncta appeared along the microtubule shaft within 2 min of incubation (Fig. 3a), consistent with selective recognition of lattice openings.

**Fig. 3.**
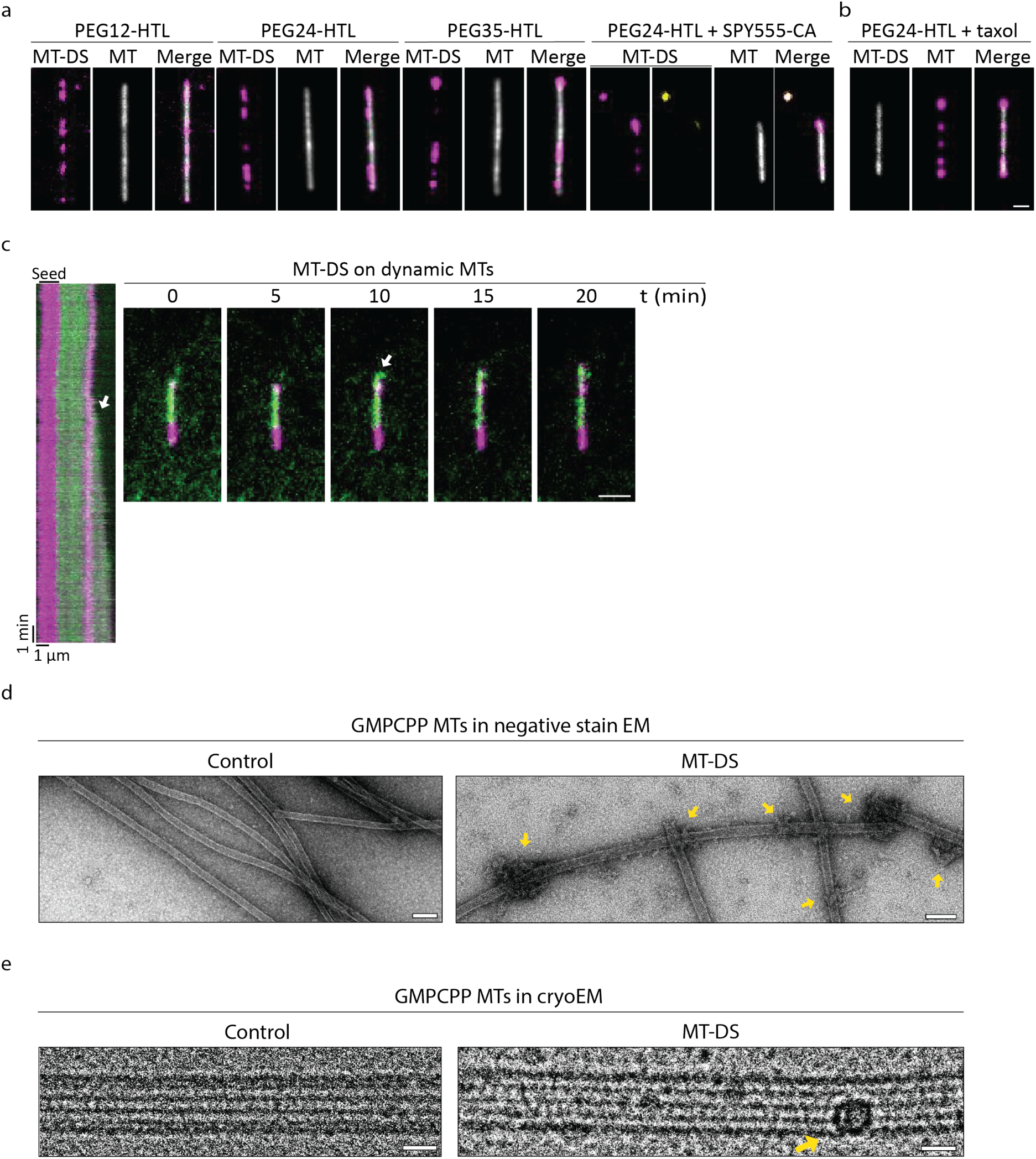
MT-DS on stable and dynamic microtubules. **a,b** Representative fluorescence images of MT-DS variants (magenta) bound to GMPCPP-microtubule (MT, white) (**a**) Labeling patterns of PEG12-HTL, PEG-24HTL, PEG35-HTL, and PEG24-HTL supplemented with SPY555-CA (yellow). (**b**) PEG-24HTL labeling of taxol-pretreated, GMPCPP-stabilized microtubules. Microtubules were pre-incubated with taxol, no taxol was added to the final solution. Scale bar: 1 um. (**c**) Kymograph and representative fluorescence images of a dynamic microtubule (green) growing from a microtubule seed (magenta) in presence of 8 nM MT-DS (magenta). Microtubule growth pauses for ∼ 10 min upon binding of MT-DS and subsequently resumes, with polymerization continuing around the probe as indicated in the kymograph by a white arrow. Scale bar: 2 μm. **d,e** Representative analysis of MT-DS-labeled microtubules (**d**) by negative stain electron microscopy (EM) and (**e**) 2D cryo-EM images GMPCPP-stabilized microtubules incubated with MT-DS or control conditions. Yellow arrows indicate MT-DS particles associated with the microtubule lattice. Scale bars, 100 nm (negative stain EM) and 15 nm (cryo-EM).

Among the tested variants, MT-DS assembled with an intermediate PEG linker length (PEG24-HTL) demonstrated bright, spatially confined signals localized to discrete lattice sites with minimal diffuse background. The shorter linker variant (PEG12-HTL) yielded comparable results, although the signal was slightly less discrete. The longer linker variant (PEG35-HTL) displayed a similar labeling pattern; however, its purification yield was substantially lower (Fig. S2b). Reduced multivalency by 50% decreased microtubule labeling efficiency (Fig. 2a), likely due to reduced affinity for open lattices. Based on these results, we selected MT-DS (PEG24-HTL) for all subsequent experiments.

### MT-DS on stabilized and dynamic microtubules

Because taxol is commonly used in microtubule assays, we next tested whether taxol interferes with MT-DS binding. Microtubules were polymerized in the presence of GMPCPP and taxol and subsequently incubated with MT-DS. MT-DS labeling was indistinguishable between taxol-pretreated GMPCPP microtubules, with no taxol in the final buffer, and microtubules prepared without taxol. These results indicate that MT-DS binding is compatible with prior taxol occupancy, likely due to the high taxol unbinding rate^24^ and is therefore compatible with standard stabilization conditions (Fig. 3b).

Because MT-DS is a multivalent microtubule-binding probe that can associate with microtubule ends, we next examined whether it perturbs microtubule dynamics. To this end, we polymerized dynamic microtubules from GMPCPP/taxol-stabilized seeds in the presence of GTP and MT-DS and monitored growth by TIRF microscopy. MT-DS localized to the growing microtubule tip during polymerization (Fig. 3c). Following MT-DS binding, microtubule growth transiently paused for approximately 10 min before resuming elongation for an additional ∼10 min. This behavior resembles that reported for compounds that block individual protofilaments, such as eribulin, where microtubules continue to polymerize around the obstruction^26^. Thus, while MT-DS slows microtubule growth kinetics, it does not irreversibly block polymerization or induce catastrophic depolymerization. Instead, microtubules are able to resume elongation after a transient pause (Fig. 3c). These observations indicate that MT-DS can be used with caution in assays involving dynamic microtubules.

To directly visualize MT-DS association with microtubules, we performed negative stain EM (Fig. 3d). While MT-DS binding to microtubule was readily detected, the underlying damage was impossible to visualize due to the electron-dense nature of the probe on the microtubule and the staining procedure. To avoid this negative-contrast artifact, we next used cryoEM. This approach improved visualization of MT-DS bound to microtubules, but the underlying damage sites themselves remained undetectable (Fig. 3e).

### MT-DS sensitively detects pre-existing lattice defects

To test whether MT-DS reports intrinsic lattice damage, we assembled GMPCPP-stabilized microtubules under polymerization conditions known to modulate lattice integrity^6,27^. As expected, microtubules assembled rapidly at high tubulin concentration and elevated temperature (medium-damage conditions) exhibited a higher density of MT-DS puncta than microtubules assembled slowly at low tubulin concentration and reduced temperature - low-damage condition (Fig. 4a).

**Fig. 4.**
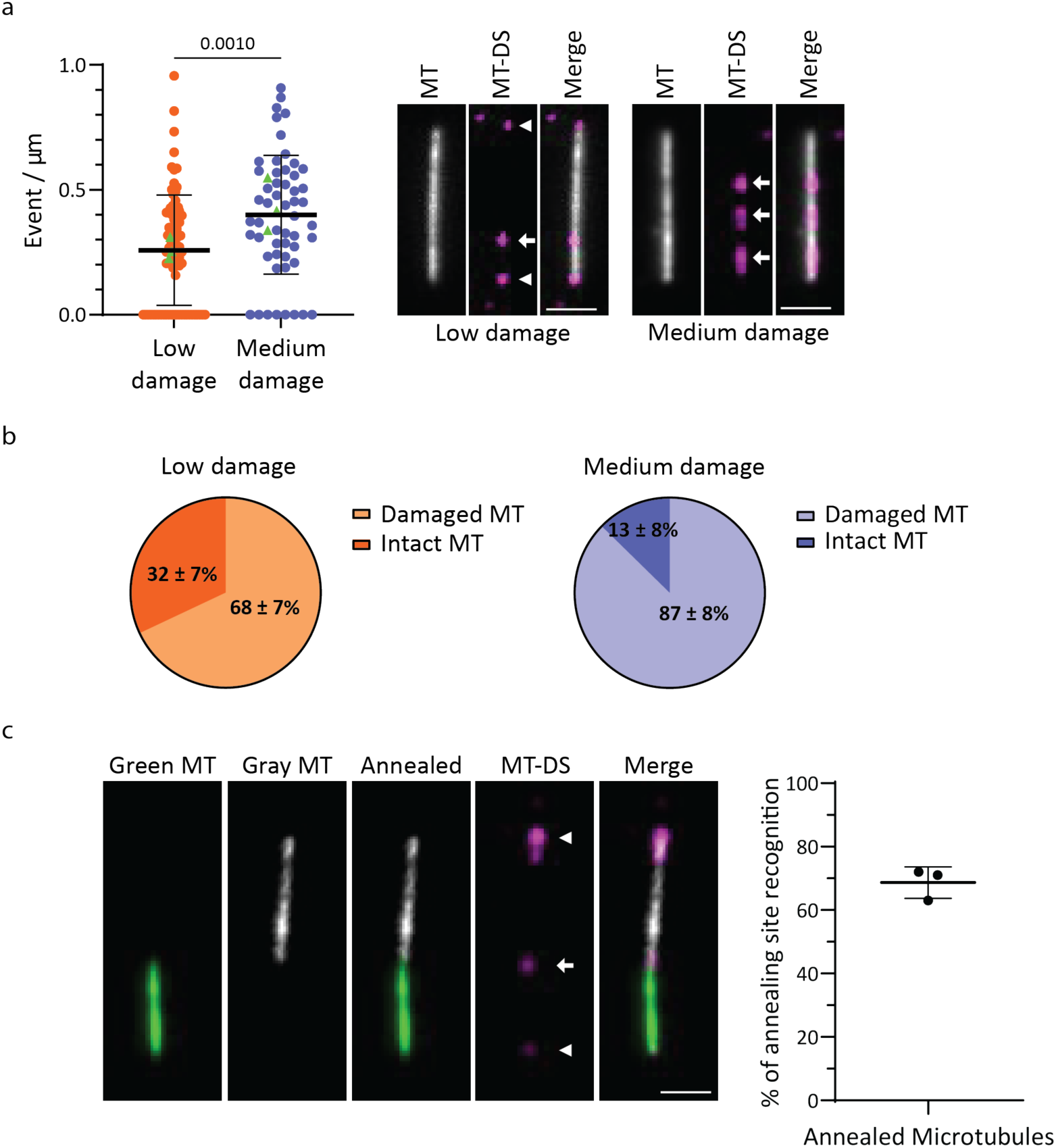
MT-DS detects differences in microtubule damage density and labels annealing sites. (**a**) Frequency of MT-DS damage recognition along GMPCPP-stabilized microtubules, end binding events were excluded from analysis. Data are shown for low-damaged (orange, n=77), and medium-damaged (blue, n=53) microtubules of three independent experiments (see Methods for more details). Replica means are indicated by green triangles. Statistics: one-way ANOVA. Mean with SD. Representative fluorescence images are shown of MT-DS (magenta) on low- or medium-damaged GMPCPP-stabilized microtubules (white). Arrow points to analyzed damage events and arrowhead to end binding. Scale bar 2 µm. (**b**) Pie charts showing the fraction of damaged or intact microtubules (MT) for low- or medium-damaged microtubules from **a**. (**c**) Representative fluorescence images of two annealed low-damaged microtubules (green and gray), with MT-DS (magenta). MT-DS labels both microtubule ends and the annealing site. Right, percentage of annealing site recognition by MT-DS for all analyzed microtubules (n=102) from three independent experiments. Scale bar 2 µm.

Quantification revealed a damage density of 0.40 events per μm under medium-damage conditions and 0.26 events per μm under low-damage conditions, comparable with previously reported results^27^. Consistently, the fraction of microtubules lacking detectable damage was significantly higher under low-damage conditions (32 ± 7%), representing an approximately 2.5-fold increase relative to medium-damage microtubules (Fig. 4b). These results demonstrate that MT-DS sensitively reports differences in lattice integrity arising from distinct assembly regimes.

To further probe MT-DS’s ability to detect localized lattice imperfections, we generated end-to-end annealing junctions by polymerizing microtubules from differently labeled tubulin pools and allowing them to anneal. Because annealing sites are prone to protofilament mismatches, they represent a defined source of structural discontinuity^28^. MT-DS robustly labeled these junctions, detecting lattice openings at approximately 70% of annealing sites (Fig. 4c). Together, these data establish MT-DS as a reliable reporter of pre-existing microtubule lattice defects.

### MT-DS outperforms repair-based assays in precision and temporal resolution

We next directly compared MT-DS to the tubulin repair assay, the current state-of-the-art method for detecting microtubule lattice damage. Repair assays exhibited high and heterogeneous background due to nonspecific tubulin interaction with the glass surface, even after extensive washing. As a result, time-resolved imaging of fluctuating microtubules is required to distinguish genuine tubulin incorporation from surface-bound aggregates that frequently exhibit higher fluorescence intensity than incorporation sites (Fig. 5a,b). In contrast, after washout, MT-DS produced high-contrast, discrete signals with negligible background, enabling unambiguous detection of damage sites without thresholding or motion analysis (Fig. 5a,b).

**Fig. 5.**
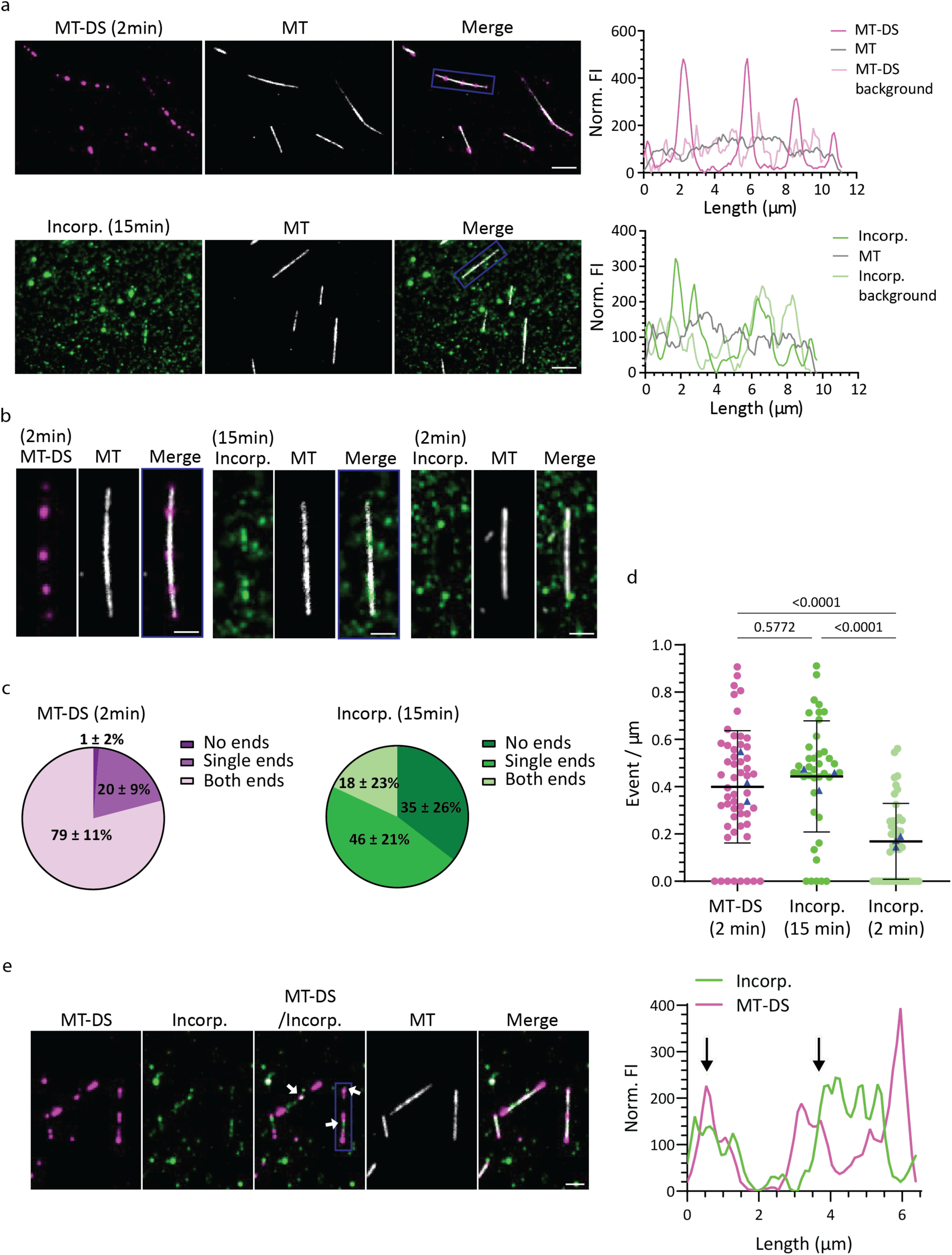
MT-DS outperforms repair assay for detecting microtubule damage. (**a**) Representative fluorescence images showing MT-DS labeling (magenta) after 2 min incubation on GMPCPP-stabilized microtubules (white) compared to incorporation of differently labeled tubulin (Incorp., green) after 15 min. Line scans of normalized fluorescence intensities (Norm. FI) are shown for microtubules in boxes, displaying the signal profiles for MT-DS or incorporated tubulin, the respective background and the microtubule (see Methods for details). Scale bars: 5 μm. (**b**) Zoom-ins of the boxed areas in **a** for 2 min MT-DS, 15 min incorporated tubulin, and representative images for 2 min incorporated tubulin. Scale bars: 2 μm. (**c**) Pie charts showing the probability (%) that MT-DS (2 min) or incorporated tubulin (15 min) detect both microtubule ends, a single end, or no ends. Mean with SD. (**d**) Frequency of damage recognition in events per μm along microtubules; end binding events were excluded from analysis. Data are shown for MT-DS (2 min, n=53), and incorporated tubulin (15 min, n=40; or 2 min, n=42) each from three independent experiments. Replica means are shown by blue triangles. Statistics: one-way ANOVA. Mean with SD. (**e**) Representative fluorescence images of GMPCPP-stabilized microtubules (white) incubated simultaneously with MT-DS (magenta) and differently labeled tubulin (Incorp., green) for 15 min. White arrows indicate colocalization events of MT-DS and incorporated tubulin. Scale bar: 2 μm. Line scans of normalized fluorescence intensities (Norm. FI) are shown for MT-DS and incorporated tubulin (Incorp.) for the microtubule in the boxed area; back arrow marks the colocalization sites.

Within 2 min, MT-DS detected both ends of GMPCPP-microtubules with a probability of 79 ± 11% (Fig. 5c). By comparison, a repair assay with 15 min of tubulin incorporation detected one microtubule end with a probability of 46 ± 21% (Fig. 5c). This limited end detection reflects the limited efficiency of microtubule polymerization from stabilized seeds^29,30^ and the depolymerization that occurs during washout of unincorporated tubulin.

Along the microtubule shaft, MT-DS detected damage at a frequency of 0.4 events per µm within 2 min, matching the efficiency of a 15-min repair assay (Fig. 5d). Reducing the tubulin incorporation time to 2 min reduced detectable events by 2.6-fold (Fig. 5d), reflecting insufficient time for lattice repair to occur. In contrast, MT-DS maintained full sensitivity, underscoring its superior temporal resolution.

Colocalization experiments confirmed that MT-DS puncta correspond to bona fide lattice defects: MT-DS signals overlapped with sites of tubulin incorporation when both assays were applied simultaneously (Fig. 5e). As we observed a reduction in overall incorporation and MT-DS fluorescence intensity - suggesting potential interference between the two approaches - we recommend using MT-DS or repair assays independently to avoid confounding effects.

### MT-DS reports kinesin-1Δ6–induced lattice damage

The kinesin-1Δ6 mutant has previously been shown to induce lattice damage on GMPCPP-stabilized microtubules. Consistent with these reports, incubation of GMPCPP-stabilized microtubules with kinesin-1Δ6 at a motor:tubulin ratio of 1:24 resulted in visible lattice distortions, including characteristic kinks along the microtubule shaft. These damages were readily detected by electron microscopy (EM) and were also robustly labeled by MT-DS using TIRF microscopy (Fig. 6a).

**Fig. 6.**
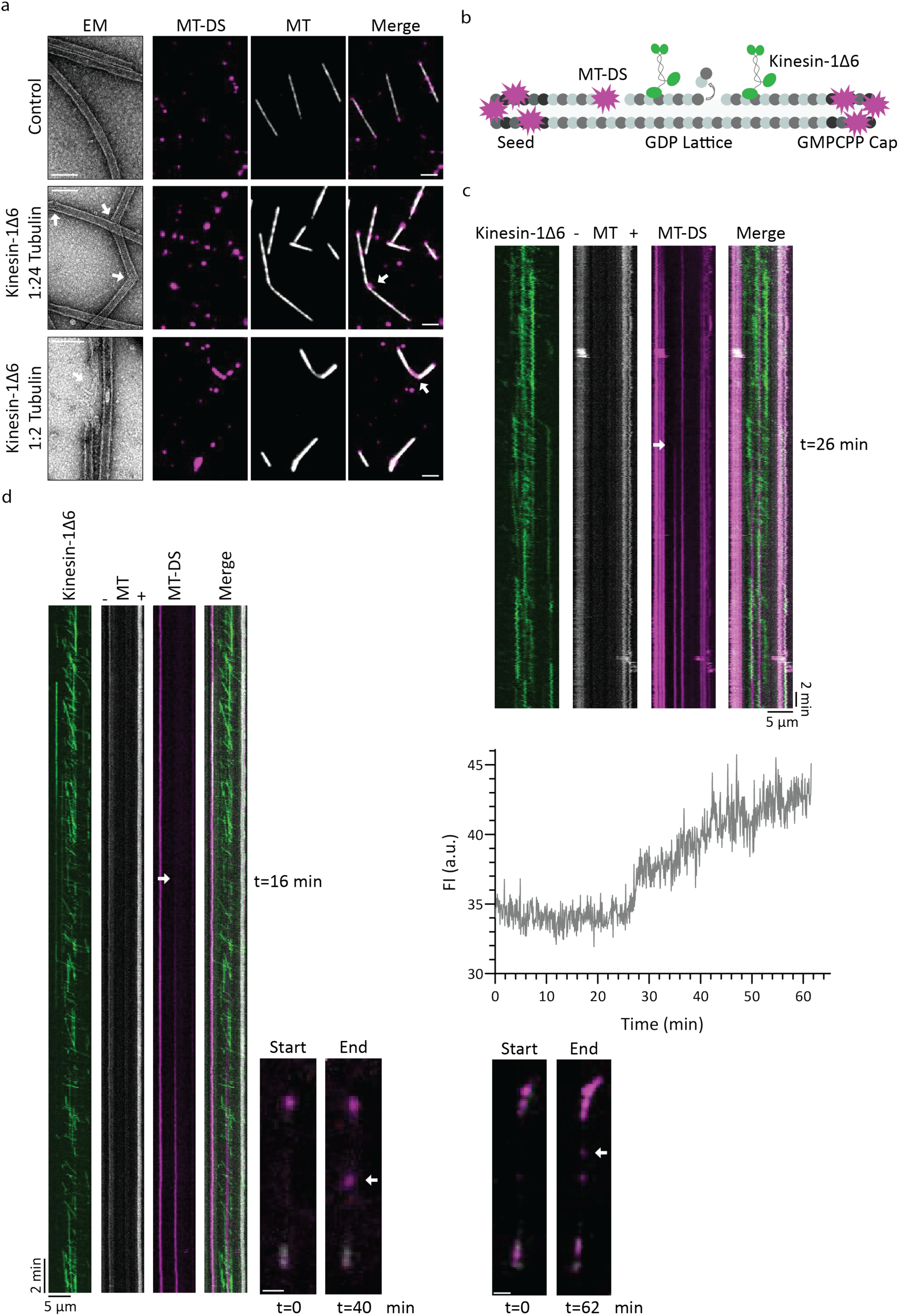
MT-DS detects kinesin-1Δ6-induced microtubule damage over time. (**a**) Electron microscope (EM) images showing damage sites. Representative fluorescence images showing detection of damage sites by MT-DS (magenta) on GMPCPP-stabilized microtubules (MT, white) in control conditions or after incubation with kinesin-1Δ6 at motor:tubulin ratios of 1:24 or 1:2. White arrows point at microtubule damage sites. Scale bars: 2 μm (fluorescence), 100 nm (EM). (**b**) Schematic of the experimental setup: Kinesin-1Δ6 motors running along GDP microtubules that are nucleated from seeds and capped with GMPCPP-tubulin. MT-DS labels damage sites and microtubule ends. **c,d** Kymographs showing kinesin-1Δ6 motility (green) along microtubules (white) together with MT-DS labeling (magenta). Fluorescence images at the start and end of each experiment are shown next to the kymographs. Scale bars: 1 μm. Imaging began with a delay of 4–10 min after addition of MT-DS and kinesin. (**c**) 1 nM kinesin-1Δ6. (**d**) 2 nM kinesin-1Δ6. White arrows indicate the onset of MT-DS accumulation at damage sites. Right, quantification of MT-DS fluorescence intensity (FI) at these sites over time.

Increasing the kinesin-1Δ6 concentration to a motor:tubulin ratio of 1:2 led to more severe lattice damage, characterized by large lattice openings and pronounced microtubule softening. These high-severity defects were again apparent by both EM and MT-DS labeling (Fig. 6a). Together, these results demonstrate that MT-DS faithfully reports kinesin-induced lattice damage across a wide range of damage severities and structural manifestations.

### MT-DS reveals de novo lattice damage during kinesin motility

An open question in the microtubule field is whether motor proteins generate lattice damage de novo or primarily amplify pre-existing lattice imperfections arising during polymerization. We used MT-DS to directly address this question by visualizing damage formation in situ during motor activity.

To minimize pre-existing lattice defects, we polymerized GDP-microtubules from GMPCPP/taxol-stabilized seeds and capped the growing ends by addition of GMPCPP-tubulin to prevent depolymerization. Under these conditions, kinesin-1Δ6 was added at low nanomolar concentrations (1 or 2 nM) together with MT-DS to the flow chamber. After assembly, the samples were mounted and imaged by time-lapse TIRF microscopy, resulting in a delay of 4-10 min between motor/MT-DS addition and start of imaging (Fig. 6b-d, see Methods for details).

At the beginning of the assay, MT-DS was already bound to structural transitions along the lattice: boundaries between seed, GDP-microtubule and GMPCPP-cap. During motor motility, MT-DS accumulated at discrete sites along the GDP lattice. In some cases, MT-DS binding was observed within minutes of motor addition, whereas in rare events MT-DS puncta emerged only after extended imaging periods, up to 25 min, indicating the progressive formation of new lattice damage sites. MT-DS fluorescence intensity at individual sites increased over time (Fig. 6e), indicating probe accumulating at a damage site.

Together, these results imply that kinesin-1Δ6 actively induces de novo microtubule lattice damage while moving along the microtubule shaft, in addition to the possibility to amplify existing defects. By enabling direct, time-resolved visualization of damage formation during motor activity, MT-DS resolves a central debate in the field and establishes motor-driven lattice damage as a dynamic and observable process.

## DISCUSSION

Understanding the structural integrity of microtubules is essential for elucidating their roles in diverse cellular processes, including intracellular transport, cell division, and cytoskeletal organization. Lattice defects can have functional consequences for microtubule resilience, such as increased mechanical fatigue under repeated bending^6^. Simulations suggest that even rare lattice defects (2%) can dramatically reduce the force required to break a microtubule^31^. Increasing evidence indicates that lattice damage and repair within the microtubule shaft are intrinsic features of microtubule dynamics^3,18^. Sites of lattice damage can undergo repair through the incorporation of free tubulin, generating GTP-tubulin islands within the shaft that promote rescue events and locally stabilize the microtubule. In this way, damage formation and repair directly influence microtubule lifetime by modulating the balance between catastrophe and rescue. Moreover, the interplay between molecular motors and the microtubule lattice involves a balance between dimer removal and self-repair mechanisms. Motor-induced damage can be compensated by tubulin incorporation, underscoring the importance of preserving an intact microtubule lattice^4^. However, direct insights into spatial distribution and dynamics of these lattice defects have remained largely inaccessible.

Current strategies to visualize microtubule damage are limited. Fluorescence-based approaches are indirect as they rely on microtubule repair through incorporation of differentially labeled tubulin. Although this strategy serves as a proxy for damage, it does not directly reveal the underlying lattice defect. EM provides essential insights into lattice ultrastructure, including local losses of tubulin density at damage sites. However, EM requires fixed samples and therefore captures only static snapshots of a dynamic process with limited throughput. As a result, the density, localization and temporal evolution of lattice damage remain poorly defined, limiting efforts to link lattice integrity to microtubule function.

The fluorescent microtubule-stabilizing agent (MSA) Fchitax-3 has been used as a visual reporter of drug-induced lattice defects, although it does not report intrinsic lattice damage. Fchitax-3 preferentially binds to plus ends in a pre-catastrophe state, producing bright accumulation “hotspots” that correspond to stabilized lattice patches (stable rescue sites). The probe accumulates at sites of protofilament number switches, where geometric mismatches generate permanent lattice defects that prevent complete lattice repair^32^. Because it labels MSA-induced protofilament transitions rather than pre-existing lattice defects, Fchitax-3 does not function as a reliable sensor of microtubule damage. In addition, custom optical severing tools with engineered severing enzymes have enabled controlled microtubule disassembly, revealing the functional consequences of mechanical breaks^33^. However, they do not directly label or quantify the location of microtubule lattice defects.

MT-DS fills this methodological gap by providing a fluorescent probe that selectively labels microtubule lattice openings in vitro. Unlike taxane-based probes such as SiR-tubulin, which uniformly labels microtubules, MT-DS specifically reports lattice openings. This capability exceeds current tools by allowing localization of defects and monitoring of their formation and dynamics. Using MT-DS, we identified intrinsic lattice defects of stabilized and dynamic in vitro-polymerized microtubules (Fig. 4). Enrichment of MT-DS at annealing sites suggests that lattice imperfections are an inherent feature of microtubule assembly (Fig. 4c). Moreover, MT-DS revealed that the kinesin-1Δ6 mutant actively generates lattice damage while running along microtubules, inducing characteristic kinks and sheet openings (Fig. 5). Over extended time frames, persistent MT-DS binding at damage sites may also influence their evolution by sterically hindering complete lattice repair. In such cases, defects could increase in size, potentially explaining the extended incorporation site partially overlapping with probe (Fig. 5e) and the progressive increase in MT-DS fluorescence intensity observed at kinesin-1–induced damage sites within the GDP lattice (Fig. 6c-d). Together, these results position lattice integrity as a directly measurable and functionally relevant layer of microtubule regulation.

Looking forward, extending damage-sensitive probes to living cells would enable direct mapping of lattice defects in response to mechanical stress, motor activity and pharmacological perturbations. Such approaches would provide a crucial link between lattice integrity and microtubule-associated processes in cellular environments, advancing our understanding of how microtubule networks are maintained under physiological and pathological conditions.

## METHODS

### Plasmid generation

To generate the plasmid for 24mer-HaloTag7 bacterial expression, the sfGFP sequence from pET29b-T33-21AB-sfGFP (a kind gift from David Baker) was replaced by the HaloTag7 sequence (plasmid a gift from Stefan Matile) using Gibson assembly. The highly damaging constitutively active kinesin-1 mutant (kinesin-1Δ6-GFP) was generated as previously described^14^ and cloned into pET17-K560-GFP-His (addgene #15219) using EcoRI/AgeI restriction enzymes.

### Protein expression and purification

For purification of the <u>24mer-HaloTag7</u>, *E.coli* BL21 (DE3 pLysS) cells were transformed and grown at 37 °C until an optical density at 600 nm of 0.6-0.8 and induced for protein expression with 0.5 mM IPTG at 18 °C overnight. Cells were harvested and lysed in lysis buffer (50 mM Tris pH 8, 500 mM NaCl) supplemented with 1 mM DTT, 1 mM PMSF and protein inhibitor cocktail (Roche, 11873580001) and subsequently sonicated. Lysed cells were cleared by centrifugation at 12000 rpm for 30 min at 4 °C. Proteins were affinity purified using a 1mL HisTrap HP column (GE Healthcare), washed in lysis buffer supplemented with 25 mM imidazole and eluted with 500 mM imidazole within 25 column volumes. Fractions were pooled and concentrated using Amicon 30K Centrifugal Filters (Millipore). Concentrated proteins were loaded onto a size-exclusion chromatography using a HiLoad 16/600 Superdex 200 column (GE Healthcare) in 25 mM Tris pH 8, 150 mM NaCl, 2mM DTT. Collected fractions were concentrated and concentration determined by Bradford assay. Proteins were snap-frozen in liquid nitrogen and stored at -80 °C until protein labeling.

For <u>kinesin-1Δ6-GFP</u> (referred to as: kinesin-1Δ6) a similar protein purification protocol was used. Protein expression was induced with 0.5 mM IPTG at 16 °C overnight. Cells were lysed in 100 mM Na_2_HPO_4_/NaH_2_PO_4_ pH 7.5, 600 mM KCl, 3 mM MgCl_2_, 10% glycerol, 20 mM β-mercaptoethanol, supplemented with 10 mM imidazole, protein inhibitor cocktail and 100 mM ATP addition right before lysis. Cell lysates were ultracentrifuged at 40000 rpm for 50 min at 4 °C. Proteins were loaded on a 1 mL HisTrap HP column were washed with 30 mM imidazole, washed again with 60 mM and eluted with a 60-500 mM imidazole gradient over 30 column volumes. Size-exclusion chromatography was performed in 10 mM HEPES pH 7.4, 150 mM KCl, 1 mM MgCl_2_, 0.05 mM ATP, 1 mM DTT and concentrated proteins were supplemented with 20% sucrose before snap-freezing and storage at -80 °C.

### 24mer-HaloTag7 labeling for MT-DS generation

The purified 24mer-HaloTag7 protein was incubated with a 2-fold molar excess of HaloTag ligand diluted in 1 mL of purification buffer (25 mM Tris pH 8, 150 mM NaCl, 2mM DTT) on a rotating wheel in the dark at 4 °C for 30 min. The synthesized HaloTag ligands PEG12-HTL, PEG24-HTL and PEG35-HTL were used, as well as SPY555-CA (Spirochrome). For the combination variant PEG24-HTL and SPY555-CA in a 10:1 molar ratio was used due to an expected higher labeling efficiency for the commercial HaloTag ligand SPY555-CA compared to PEG24-HTL which resulted in a 50% labeling ratio (Fig. S2b). After incubation, the labeled 24mer-HaloTag7 variants were purified by size-exclusion chromatography individually to remove excess unbound ligand. Collected fractions were concentrated and concentration determined by Bradford assay. Labeled proteins were supplemented with 10 % glycerol, aliquoted, snap-frozen in liquid nitrogen and stored at -80 °C.

### Tubulin purification and labeling

Tubulin was purified from bovine brain by two polymerization/depolymerization cycles in High-Molarity PIPES buffer as described previously^34^. In brief, a first polymerization/depolymerization was performed in High-Molarity PIPES buffer (1 M PIPES pH 6.9, 10 mM MgCl_2_, 20 mM EGTA, 1.5 mM ATP, 0.5 mM GTP) supplemented with 1:1 glycerol and Depolymerization buffer (50 mM MES at pH 6.6, 1 mM CaCl_2_) respectively. A second polymerization/depolymerization was performed with polymerization in High-Molarity PIPES buffer and depolymerization in 0.25x BRRB80 with addition of 5x BRB80 after 15 min to reach 1xBRB80 (80 mM PIPES at pH 6.8, 1 mM MgCl_2_, 1 mM EGTA).

Purified tubulin was labeled with ATTO-488, ATTO-565, ATTO-647 (ATTO-TEC GmbH) or biotinylated tubulin, adapted from a previously published protocol^35^. Briefly, tubulin was polymerized in glycerol PB solution (80 mM PIPES pH 6.8, 5 mM MgCl2, 1 mM EGTA, 1 mM GTP, 33% glycerol) for 30 min at 37 °C and placed in layers onto cushions of 0.1 M NaHEPES pH 8.6, 1 mM MgCl_2_, 1 mM EGTA and 60% glycerol and centrifuged. The pellet was resuspended in resuspension buffer (0.1 M NaHEPES pH 8.6, 1 mM MgCl_2_, 1 mM EGTA and 40% glycerol) followed by incubation for 10 min at 37 °C with either 1/10 volume of 100 mM ATTO-488, -565, or -647-NHS-fluorochromes or for 20 min at 37 °C with 2 mM NHS-biotin. Labeled tubulin was sedimented onto cushions of 1x BRB80 supplemented with 60 % glycerol, resuspended in BRB, and a second polymerization/depolymerization cycle was performed. Labeled and unlabeled tubulin was aliquoted and stored in liquid nitrogen.

### Microtubule assembly

GMPCPP-stabilized microtubules (medium-damaged) were assembled by mixing 10% ATTO-488 or ATTO-565 labeled tubulin with 60% unlabeled and 30% biotinylated tubulin with a final concentration of 5 μM in 1x BRB80 buffer (80 mM PIPES pH 6.8, 1 mM MgCl_2_, 1 mM EGTA) supplemented with 0.5 mM GMPCPP. The mixture was incubated for 20 min at 37 °C, followed by two subsequent additions of 1 μM tubulin for 10 min at 37 °C, centrifuged at 12000 rpm for 15 min at RT and microtubules were resuspended in 1x BRB80. Note that this procedure was used when the text refers to GMPCPP-stabilized microtubules, if not stated otherwise.

For assembling low-damaged GMPCPP-stabilized microtubules, 2.5 μM of 10% ATTO-488 or ATTO-565 labeled tubulin with 60% unlabeled and 30% biotinylated tubulin in 1x BRB80 buffer supplemented with 0.5 mM GMPCPP were incubated for 5h at 28°C, according to Reuther et al^27^. Microtubules were centrifuged at 12000 rpm for 15 min at RT and resuspended in 1x BRB80.

### Microtubule seed preparation

Short microtubules seeds were prepared by mixing 30% ATTO-647 or ATTO-565 labeled tubulin with 70% biotinylated tubulin with a final concentration of 10 μM in 1x BRB80. The solution was supplemented with 0.5 mM GMPCPP and incubated for 45 min at 37 °C. After the addition of 1 μM paclitaxel (taxol, Sigma-Aldrich, T1912) the solution was incubated again for 30 min at 37 °C and subsequently centrifuged for 15 min at 14000 rpm at room temperature (RT). The microtubule pellet was resuspended in 1x BRB80 supplemented with 0.5 mM GMPCPP and 1 μM taxol. Seeds were aliquoted and stored in liquid nitrogen.

### Flow chamber preparation

Glass slides and Coverslips were cleaned in two steps: sonication in 1 M NaOH for 40 min, rinsing in bidistilled water, followed by sonication in 96% ethanol for 40 min and repeated rinsing in bidistilled water. Slides and coverslips were dried with an air gun and plasma-treated in a plasma cleaner (Electronic Diener, Plasma surface technology). Slides were incubated for two days in tri-ethoxy-silane-PEG (1mg/mL in 96% ethanol and 0.02% HCl, Creative PEGWorks) and coverslips in a 1:5 mixture of tri-ethoxy-silane-PEG-biotin with tri-ethoxy-silane-PEG, for two days with gentle agitation at RT. Slides and coverslips were washed in 96% ethanol and bidistilled water, dried with an air gun and stored in a sealed container at 4 °C. Flow chambers were assembled by fixing a piece of treated coverslip on a treated slide by using double-sided tape. The chamber holds around 10 μL.

### Microtubule repair assay

The repair assay was adapted from the recently published protocol^36^. In a flow chamber, 30 μL neutravidin (50 μg/mL, Thermo Fisher Scientific, 31000) were added and incubated for 1 min. After washing with 1x BRB80 (95 μL), GMPCPP-microtubules diluted in 1x BRB80 (95 μL) were added and incubated for 2 min. Unattached microtubules were washed off twice with 1x BRB80 (95 μL each) and then 5 μM ATTO-488 or ATTO-565 labeled tubulin in 1x BRB80 supplemented with 1 mM GTP were added to the flow chamber. The flow chamber was incubated in a closed wet chamber for 2 or 15 min at 37 °C for tubulin incorporation. The excess of free tubulin was washed off twice with 1x BRB80 (95 μL each) and the chamber sealed for immediate microscopy analysis.

### Microtubule damage labeling over time with SiR-CabazyLys-Prop or D40/D70-PEG3-SiR-CabazyLys

GMPCPP-stabilized microtubules were added to in-house prepared microfluidic flow chambers with 2×100 mm channels, as described for regular flow chambers. After washing off microtubule excess, the chamber was placed inside the microscopy holder and image acquisition was started. While imaging, a solution of 10nM SiR-CabazyLys-Prop, D40- or D70-PEG3-SiR-CabazyLys diluted in 1x BRB were added to the flow chamber and while images were acquired every second.

### Microtubule damage labeling with MT-DS

GMPCPP-stabilized microtubules were added to the flow chamber (as explained above). After washing off microtubule excess twice, MT-DS (50-125 nM) was added to chamber and incubated for 2 min at RT, shorter labeling time of 1 min gave similar results. Probe excess was washed off twice with 1x BRB80 (95 μL each) and the chamber sealed for immediate microscopy analysis. Labeling of all MT-DS variants HTL13, HTL25, HTL38 and HTL25 + SPY555-CA was performed the same way. For assessing MT-DS HTL25 in the presence of taxol, GMPCPP-microtubules were polymerized as described above with a final resuspension in 1x BRB80 containing 10 μM taxol. These taxol GMPCPP-microtubules were added to the flow chamber. Note that we did not add taxol to the flow chamber.

For co-labeling of MT-DS with incorporated tubulin, a microtubule repair assay was performed as described and MT-DS (25 nM) was added to the incorporation solution for 15 min. Probe and tubulin excess was washed off twice with 1x BRB80 (95 μL each) and the chamber sealed for immediate microscopy analysis.

### Microtubule annealing

Low-damaged GMPCPP-stabilized microtubules were polymerized with either ATTO-488 or ATTO-565 labeled tubulin, as described above. Both microtubule preparations were mixed in a 1:1 ratio and incubated for 1h at 30°C for microtubule annealing. MT-DS labeling was performed as described.

### Microtubule dynamic assay

After addition of neutravidin to the flow chamber (as described above) microtubule ATTO-647 seeds diluted in 1xBRB80 (95 μL) were added, incubated for 2 min and excess washed off with 1x BRB (95 μL). A 30 μL solution containing 10 μM 10% ATTO-488 labeled and 90% unlabeled tubulin in anti-bleaching buffer [1x BRB80, 1 mM GTP, 10 mM DTT, 0.3 mg/mL glucose, 0.1 mg/mL glucose oxidase, 0.02 mg/mL catalase, 0.125% methyl cellulose (1,500 cP, Sigma-Aldrich, M0387)] with MT-DS (8 nM) was added to the flow chamber and the chamber sealed for microscopy analysis. Each position was recorded over 240 frames with an interval of 5 seconds.

### Microtubule damage generation with kinesin-1Δ6

For damaging GMPCPP-stabilized microtubules with kinesin-1Δ6, microtubules (3.5 μM tubulin) were incubated with either 1.75 μM or 84 nM of kinesin-1Δ6 for a 1:2 or 1:24 tubulin ratio respectively, in 1x BRB supplemented with 2.7 mM ATP. Control conditions were performed in 1x BRB with 2.7 mM ATP only. Reactions were incubated for 15 min at 37 °C and immediately used for further assessment. For fluorescence-based analysis, microtubules were added to the chamber and labeled with MT-DS as described above. For electron microscopy analysis, microtubules were fixed immediately as described below.

For visualization of kinesin-1Δ6-induced damage at the microscope, microtubules were polymerized from ATTO-565 seeds in a flow chamber (as described above). After washing off excess seeds, 30 μL casein (0.5 mg/mL, Sigma-Aldrich, S0406) was perfused through the chamber to reduce unspecific binding and then washed off with 1x BRB80 (95 μL). A 20 μL solution containing 10 μM 10% ATTO-565 labeled and 90% unlabeled tubulin in 1x BRB80 supplemented with 1 mM GTP and 0.1 mg/mL casein was added to the flow chamber and incubated in a closed wet chamber for 20 min at 37 °C for microtubule to polymerize from seeds. A prewarmed capping solution (20 μL) of 5 μM 70% ATTO-565 labeled tubulin and 30% unlabeled tubulin in 1x BRB80 supplemented with 0.25 mM GMPCPP and 0.1 mg/mL casein was perfused through the flow chamber to exchange the polymerization buffer and incubated for 15 min at 37 °C for microtubule capping. An imaging solution (20 μL) containing 1 or 2 nM kinesin-1Δ6-GFP, 2.7 mM ATP, 50 mM KCl, 0.1 mg/mL casein, 8 nM MT-DS, 1 μM unlabeled tubulin in anti-bleaching buffer was added and the chamber sealed for microscopy analysis. Note, that between addition of MT-DS probe, kinesin-1Δ6-GFP and starting of imaging we had a time delay of about 4-10 min. This time delay results from chamber sealing, mounting on the microscope and selection of field of view. Imaging of 1 nM kinesin-1Δ6-GFP was performed over 817 frames with an interval of 4.53 seconds. The position for 2 nM kinesin-1Δ6-GFP was recorded over 1456 frames with an interval of 1.65 seconds.

### Slow microtubule lattice disassembly assays

GMPCPP-stabilized microtubules were added to a flow chamber (as described above). No anti-bleaching buffer and no free tubulin was added, and microtubule breakage was recorded over 26 frames with an interval of 4 seconds.

### SDS-PAGE and in-gel fluorescence

Samples were diluted in 1x sample buffer (60 mM Tris, 2% SDS, 10% glycerol, 5% β-mercaptoethanol, 0.005% bromophenol blue) and run on 4-20% Mini-PROTEAN® TGX Stain-Free™ Protein Gels (BIO-RAD) in 1x running buffer (25 mM Tris, 192 mM Glycine, 0.1% SDS, pH 8.3). Protein load was detected by UV exposure and fluorescence emission was detected by gel excitation (564 or 640nm) with a Fusion FX Vilber (Witec AG). Fluorescence intensity values were quantified and divided by the protein load for comparison.

### Negative stain transmission electron microscopy

For negative stain transmission electron microscopy (negative stain EM), damaged microtubules were diluted in 1x BRB to a final concentration of 3.5 µM. MT-DS (125 nM) was added and the mix incubated for 2 min. The sample was centrifuged at 12000 rpm for 10min to remove excess MT-DS and resuspended in 1x BRB. A 4 µL drop of the sample was applied directly onto glow-discharged copper EM grids (Electron Microscopy Sciences). After a one-minute incubation, the excess sample was blotted off, and the grids were washed twice with 10 µL drops of 2% uranyl acetate (Electron Microscopy Sciences), each time followed by immediate blotting. The grids were then stained with a 10 µL drop of 2% uranyl acetate for one minute, blotted, and air-dried for two minutes. Prepared grids were stored in EM grid boxes at RT until imaging. Imaging was performed using a Talos L120C transmission electron microscope operated at 120 kV and data was recorded using EPU software (ThermoFisher Scientific).

### Cryo electron tomography

Cryo-electron tomography (Cryo-ET) was performed as previously described^37^. For cryo-ET and 2D micrographs, damaged microtubules with and without MT-DS were prepared following the same procedure as for negative stain EM. A 4 µL drop of sample was applied directly onto glow-discharged Lacy Carbon, 300 Mesh, Gold grids (Electron Microscopy Services). After one-minute incubation at 37 °C and 95 % humidity, samples were automatically back-blotted and plunged immediately into liquid ethane using an EM GP2 automatic plunge freezer (Leica). Samples were stored in liquid nitrogen until imaging. All 2D micrographs and tomograms were acquired on a Talos Arctica electron microscope operated at 200kV, equipped with a Falcon 4i Direct Electron Detector and Selectris X energy filter (ThermoFisher Scientific). Data was recorded using Tomography 5.10 software (ThermoFisher Scientific). The pixel size was 1.5 Å, and the tilts were ± 60 °C with 3° between each tilt.

### Image and statistical analysis

Microscopy images were taken with an Axio Observer Inverted TIRF microscope (Zeiss, 3i) and a Prime 95B (Photometrics). A 100x objective (Zeiss, Plan-Apochromat 100x/1.46 oil DIC (UV) VIS-IR) was used. SlideBook 6X64 software (version 6.0.4) was used for image acquisition. Microscope stage conditions were controlled with a Tokai hit stage top incubator (STXG-SP400NX-SET) with humidifier (HUMID2ST). Images were analyzed using ImageJ.

For quantification of <u>MT-DS labeling and tubulin incorporation signals</u>, individual puncta along the microtubule lattice were counted, independent of the puncta size. The <u>increase in fluorescence intensity</u> <u>over time</u> was determined by measuring the probe signal (SiR-CabazyLys-Prop or dextrane-coupled) every second in 1 µm length in the middle of a single microtubule. The background intensity of 1 µm length was measured every second and subtracted from the respective data set and fluorescence intensity values were plotted over time. For the <u>line profile</u> of MT-DS or incorporated tubulin, the fluorescence intensity signals of MT-DS, incorporated tubulin and microtubules in the highlighted boxes were plotted. The background intensity was measured with the same box in the direct vicinity of the microtubule. Each line plot was normalized to the smallest value of the respective data set. The <u>probability of microtubule end</u> <u>detection</u> was determined by manually counting the number of detected ends per microtubule. The sum for no, single or both ends was divided by the total number of analyzed microtubules and values plotted as percentage with standard deviation. The <u>density of events</u> was analyzed by manually counting the number of MT-DS or incorporation signals along the microtubule lattice. The signals detected at microtubule ends were subtracted and the final event count was divided by the length of the individual microtubule. The event density for individual microtubules was plotted. The <u>percentages of damaged</u> <u>microtubules</u> were calculated by dividing the number of intact or damaged microtubules by the total number of microtubules. The <u>likelihood of annealing site recognition</u> was determined by counting the number of annealing sites with MT-DS labeling and the number of sites without labeling, divided by the total number of annealed microtubules analyzed and plotted as percentage with standard deviation. The <u>increase in fluorescence intensity for MT-DS labeling</u> along in situ-damaged microtubules was plotted over time. The background was subtracted, a linear fit performed and the coefficient of determination (R^2^) calculated.

Statistical analyses were performed by one-way ANOVA with Tukey’s multiple comparisons test. using GraphPad Prism software v.10.4.1. P values less than 0.05 were considered statistically significant. Protein structure overlays (Fig. 2) were generated using UCSF Chimera, production version 1.19 (build 42556)^38^.

## DATA AVAILABILITY

All data associated with this study are presented in the manuscript in main figures and supplementary information.

## ACKNOWLEDGEMENTS

CE has been supported by the SNSF, (TMSGI3_211433), LR, AS by the European Research Council (ERC) under the European Union’s Horizon 2020 research and innovation programme (grant agreement n° 948750, DestCilia, to SH). JT has been supported by the DIP of the Canton of Geneva. CA has been supported by the DIP of the Canton of Geneva, SNSF (310030_212563, CRSII5_216597 and TMSGI3_211433). SH has been supported by the University of Geneva, the Swiss National Science Foundation (project grants 189246 and 10000608) and the European Research Council (ERC) under the European Union’s Horizon 2020 research and innovation programme (grant agreement n° 948750, DestCilia).

## AUTHOR CONTRIBUTIONS

CE and CA conceptualized the study and designed the experiments. LR and AS synthesized the molecules used to generate the probe. MCV generated the plasmids used in this study. JT performed electron microscopy. CE performed and analyzed all experiments. SH supervised AS, CA supervised CE, JT and MCV. CE and CA wrote the manuscript. SH, LR, AS, JT edited the manuscript.

## LIMITATIONS OF THE STUDY

Because underlying microtubule damage is not directly visible in cryoEM, the minimal defect size detectable by MT-DS cannot be determined.

---

**Fig. S1.**
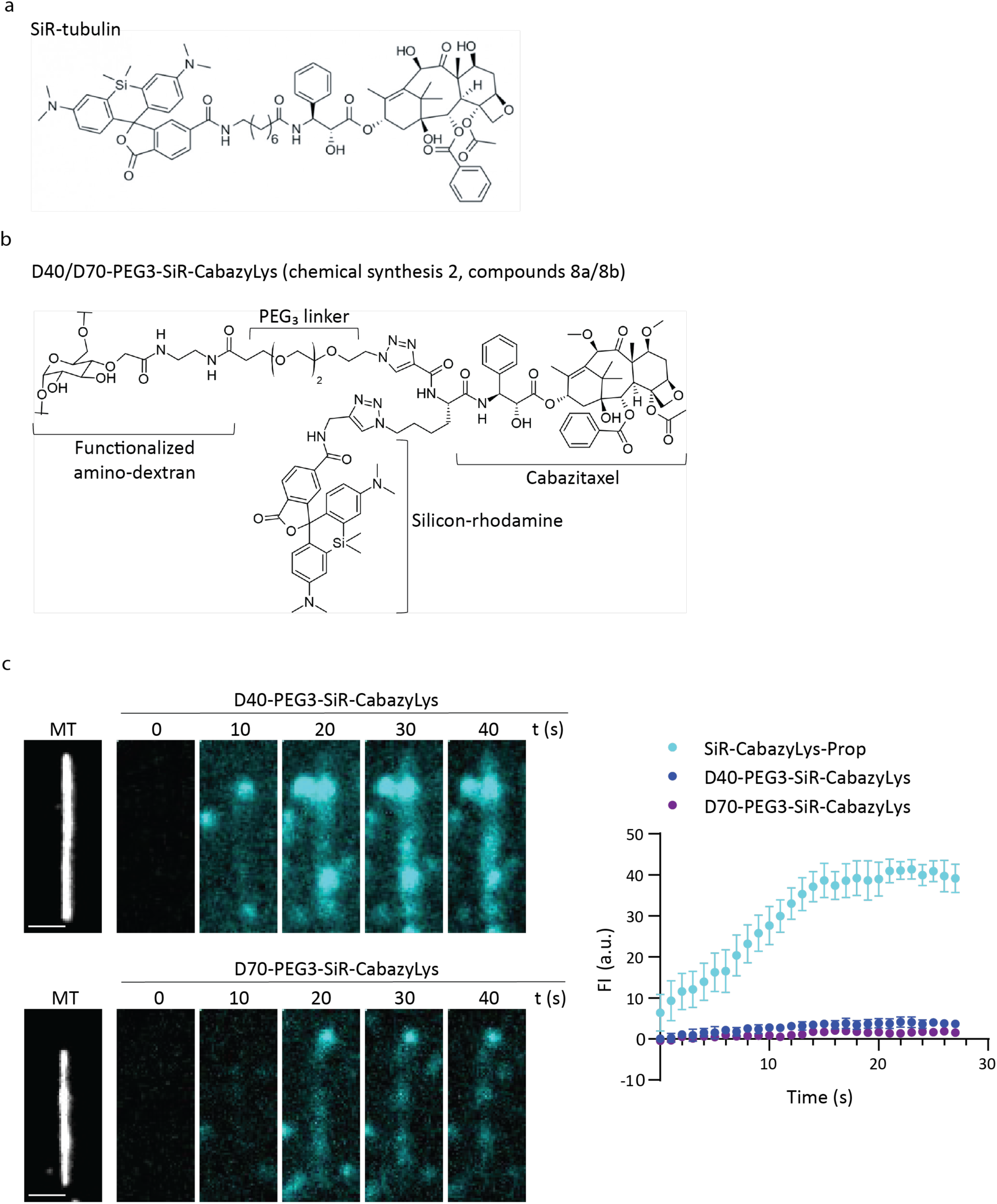
Generation of dextran-based microtubule damage probes. **a,b** Chemical structures of (**a**) SiR-tubulin^22^, and (**b**) D40/D70-PEG3-SiR-CabazyLys (chemical synthesis 2, compounds 8a/8b). (**c**) Representative fluorescence images of a GMPCPP-stabilized microtubule (MT, white) which shows microtubule labeling upon addition of D40- or D70-PEG3-SiR-CabazyLys (10 nM, cyan, auto-contrast) over time. Scale bar: 2 μm. Right, quantification of fluorescence intensity along the microtubule over time from microfluidics-based, high–temporal resolution labeling experiments following addition of SiR-CabazyLys-Prop (n = 5), D40-PEG3-SiR-CabazyLys (n = 5), or D70-PEG3-SiR-CabazyLys (n = 5).

**Fig. S2.**
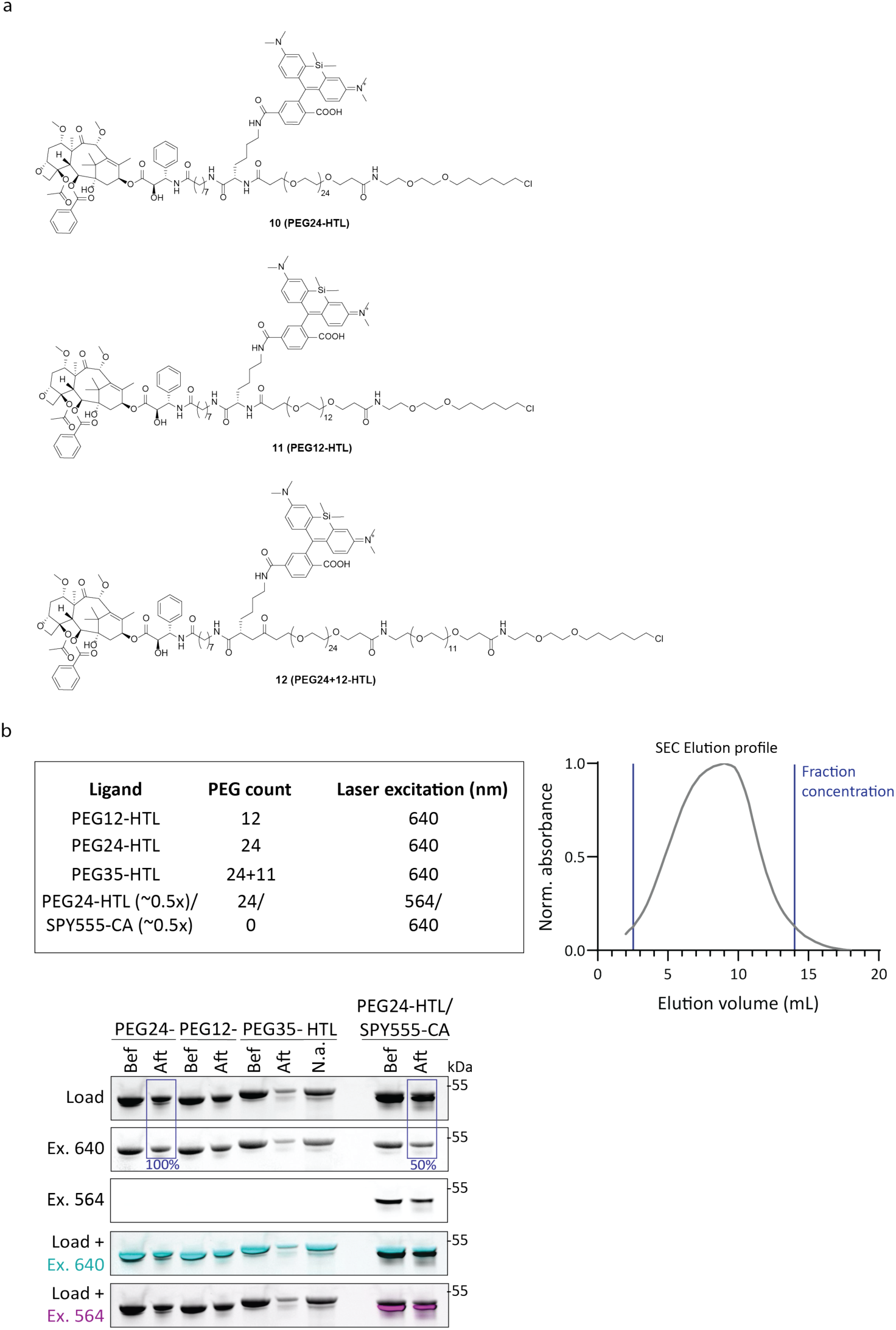
Generation and characterization of MT-DS variants. (**a**) Chemical structure of PEG24-HTL, PEG12-HTL and PEG35-HTL (chemical synthesis 1, compounds 10-12). (**b**) MT-DS variants differ in PEG linker size from 12-35 additional PEG units per HaloTag (24 tags per multimer), and by optional labeling with the HaloTag ligand SPY555-CA. Representative size exclusion chromatography (SEC) profile for the purification of each MT-DS variant are shown (right). Normalized (Norm.) absorbance is plotted as a function of elution volume; fractions collected for concentration are indicated by blue lines. Below, in-gel analysis shows protein loading and fluorescence emission signals following excitation at 640 nm (Ex. 640) or 564 nm (Ex. 564) for each MT-DS variant before (bef) and after (aft) SEC purification. Percentages highlighted in blue boxes indicate the ratio of 640 nm fluorescence signal to protein load for the indicated MT-DS variants.

